# Genome-wide and gene expression analyses of the hagfish olfactory receptors illuminate lineage-specific diversification of olfaction in basal vertebrates

**DOI:** 10.1101/2025.07.23.666228

**Authors:** Hirofumi Kariyayama, Ooi Yusuke, Hiromu Kashima, Taiki Nakanowatari, Riho Harada, Yoko Yamaguchi, Daichi G. Suzuki

## Abstract

Olfactory receptors (ORs) of vertebrates have undergone remarkable diversification, but the ancestral repertoire remains incompletely understood. Here, we present a comprehensive genome-wide survey and gene expression analyses of OR genes in hagfish, a cyclostome group that typically inhabits the deep sea with a well-developed olfactory system. We identified 48 main olfactory receptors (mORs), two vomeronasal type 1 receptors (V1Rs), 135 typical vomeronasal type 2 receptors (V2Rs), and no trace amine-associated receptors (TAARs) in the inshore hagfish (*Eptatretus burgeri*). Notably, V2Rs were highly clustered on a single scaffold, suggesting tandem gene duplications as the main mechanism of diversification. Transcriptome analysis confirmed that most of these OR genes are expressed predominantly in the olfactory organ. Using *in situ* hybridization, we further confirmed that at least some of these genes are expressed in olfactory receptor cells. These gene expression analyses support the role of these genes as functional ORs. Our findings challenge the prevailing view that typical olfactory V2Rs originated in gnathostomes and instead suggest that they were present in the common ancestor of all vertebrates but were lost in lampreys.

The extensive diversification of V2Rs in hagfish likely reflects adaptations to life in low-light and chemically complex environments. This study highlights the evolution of vertebrate sensory systems in a mosaic manner, underscoring the importance of hagfish as a key animal for reconstructing the evolutionary history of early vertebrates.

**Graphical abstract:** 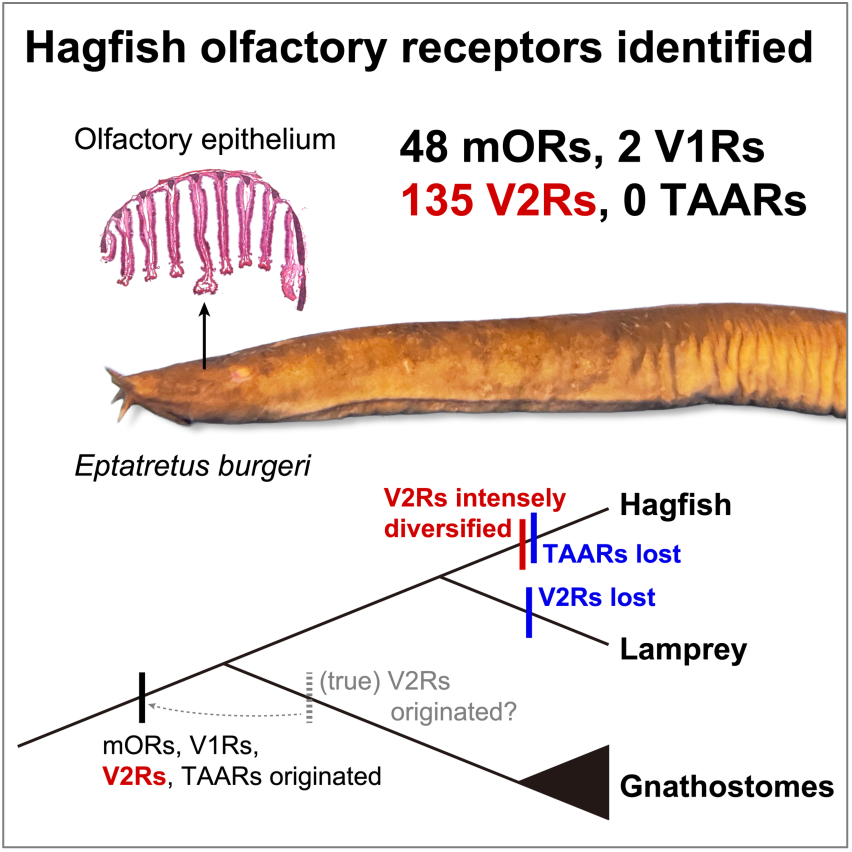

**Highlights:**

- Genomic and expression analyses reveal hagfish olfactory receptor repertoire.
- V2Rs are extensively diversified in hagfish.
- The olfactory function of V2Rs is traced back to the common ancestor of vertebrates.
- The olfactory receptor repertoires of basal vertebrates underwent mosaic evolution.

## Introduction

Olfaction is a vital sense modality for animals. In particular, it plays an even more important role in darkness because visual information is limited in such environments. It is suggested that our mammalian ancestors experienced nocturnal life, in which they heavily relied on olfaction, accompanied by enlargement of the olfactory bulb and cerebral cortex (Rowe et al. 2011). Along with this brain evolution, their olfactory receptors diversified explosively (Hughes et al. 2018).

A similar evolutionary trend may have occurred to hagfish, which belongs to the basal-most lineage of vertebrates (i.e., the cyclostomes). Hagfish typically live in light-decaying deep water and have a degenerate visual system (Fernholm and Holmberg 1975). Instead, these animals possess a highly developed olfactory bulb and telencephalon (the brain region homologous to the mammalian cerebrum), suggesting that their capacity for olfaction is correspondingly elaborated (Suzuki 2021). However, the repertoire of hagfish olfactory receptors remains largely unknown.

Apart from such habitat-specific diversification of olfaction, hagfish may provide crucial information for the early evolution of the vertebrate olfactory receptors (ORs). Within vertebrates, there are four major OR families: main olfactory receptors (mORs, or ORs in the narrow sense), type 1 vomeronasal receptors (V1Rs), type 2 vomeronasal receptors (V2Rs), and trace amine-associated receptors (TAARs). Previous research has shown that the lamprey genome contains 27 mORs (*Petromyzon marinus*; Libants et al. 2009), six V1Rs (*P. marinus* and *Lethenteron camtschaticum*; Kowatschew and Korsching 2022), one (*P. marinus*) or two (*L. camtschaticum*) V2R-like proteins, and 51 (*P. marinus*) or 32 (*L. camtschaticum*) TAARs (Dieris et al. 2021). However, Kowatschew and Korsching (2022) report that the *P. marinus* V2R-like gene is not expressed in the olfactory epithelium. Bi et al. (2021) identified more than 50 V2R-like sequences in hagfish. Still, Zhang et al. (2022) suggest that no typical V2Rs exist in extant agnathans (i.e., lampreys and hagfish) because they did not form a clade with known V2Rs in their phylogenetic analysis. To understand the ancestral repertoire of vertebrate ORs, it is therefore necessary to investigate hagfish ORs in a more thorough and detailed manner.

In this study, we conducted a comprehensive genome-wide survey and gene expression analyses of hagfish ORs. We first identified 48 mOR, two V1R, 137 V2R, but no TAAR candidate sequences in the inshore hagfish (*Eptatretus burgeri*) genome. Then, phylogenetic analysis revealed that all mOR candidates are type 1 mOR genes and that the two V1R candidates are also true vertebrate V1Rs. Regarding V2Rs, 135 among 137candidates were estimated to be typical V2Rs. We next explored the genomic distribution of the identified OR genes and found that phylogenetically close genes tend to be clustered in the genome. Transcriptome analysis indicated that most of these ORs are expressed in the olfactory organ. Using *in situ* hybridization, we further confirmed that at least some of these genes are expressed in olfactory receptor cells. These results reveal the comprehensive olfactory receptor repertoire of hagfish, suggesting that the common ancestor of vertebrates had functional olfactory V2Rs and that both lampreys and hagfish have experienced lineage-specific diversification of olfactory receptors.

## Results

### Gene identification and phylogenetic analysis of the hagfish ORs

#### mORs

In osteichthyes, including tetrapods, mORs are the biggest gene family among olfactory receptors (see Table 1). We explored hagfish gene models and genome data, and then found 48 mOR candidates (for the list of the identified hagfish OR genes, including these mOR candidates, see Supplementary File 1). We also identified 62 lamprey mOR genes that are predicted to be functional from the newest version (Timoshevskaya et al., 2023) of the sea lamprey genome (for the list of the lamprey OR genes identified in this study, see Supplementary File 2). Phylogenetic analysis revealed that all of these hagfish genes form two groups, both belonging to the type 1 mOR subfamily (Fig. 1A). Each group contained hagfish genes alone, showing monophyly with a high support value (100%). One of them (hagfish Group A) included 37 genes, belonging to a larger group with a monophyletic lamprey clade (lamprey Group A) supported by a high bootstrap value. The other (hagfish Group B) consisted of 11 genes and formed a monophyletic group with a sea lamprey gene, with a relatively low support value. No type 2 genes were identified from hagfish, although some lamprey genes were included in this subfamily as previous research suggested (Niimura 2009).

**Figure 1.**
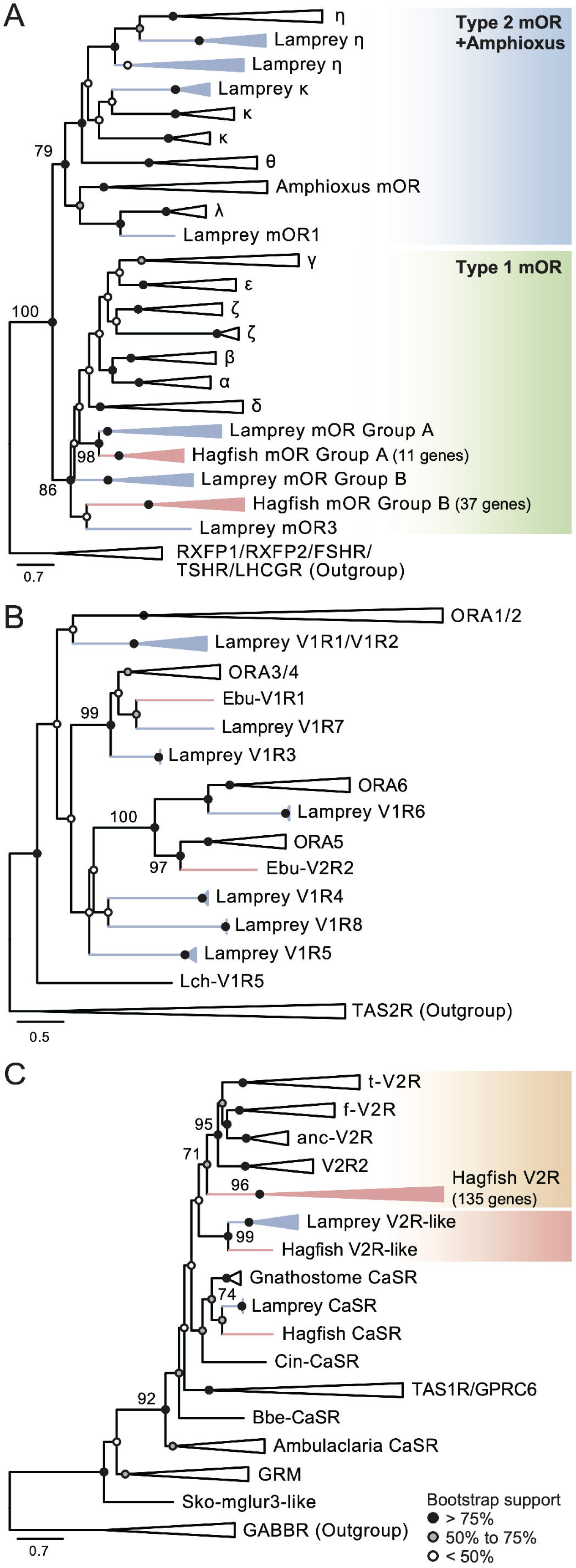
Phylogenetic analysis of mOR, V1R, and V2R genes. (A) Phylogenetic tree of mORs. White triangles indicate clades of type 1 mOR subgroups of gnathostomes, type 2 mOR subgroups of gnathostomes, amphioxus mORs, and outgroup genes. (B) Phylogenetic tree of V1R. White triangles indicate subgroups of gnathostome V1R genes and outgroup genes. (C) Phylogenetic tree of V2R. White triangles indicate subgroups of gnathostome V2R genes, gnathostome CaSR genes, gnathostome TAS1R/GPRC6 genes, Ambulacraria CaSR genes, GRMs, and outgroup genes. For all trees, red triangles and edges denote clades of hagfish genes, while blue triangles and edges mark clades of lamprey genes. The range of bootstrap support values is shown as the circles at each node (black > 75, 75 ≥ gray ≥ 50, white < 50). The numbers described on some nodes indicate exact bootstrap values. Scale bar represents the number of amino acid substitutions per site. For the list of genes used, see Supplementary File 3. For more detailed trees, see Supplementary File 4–6.

**Table 1.**
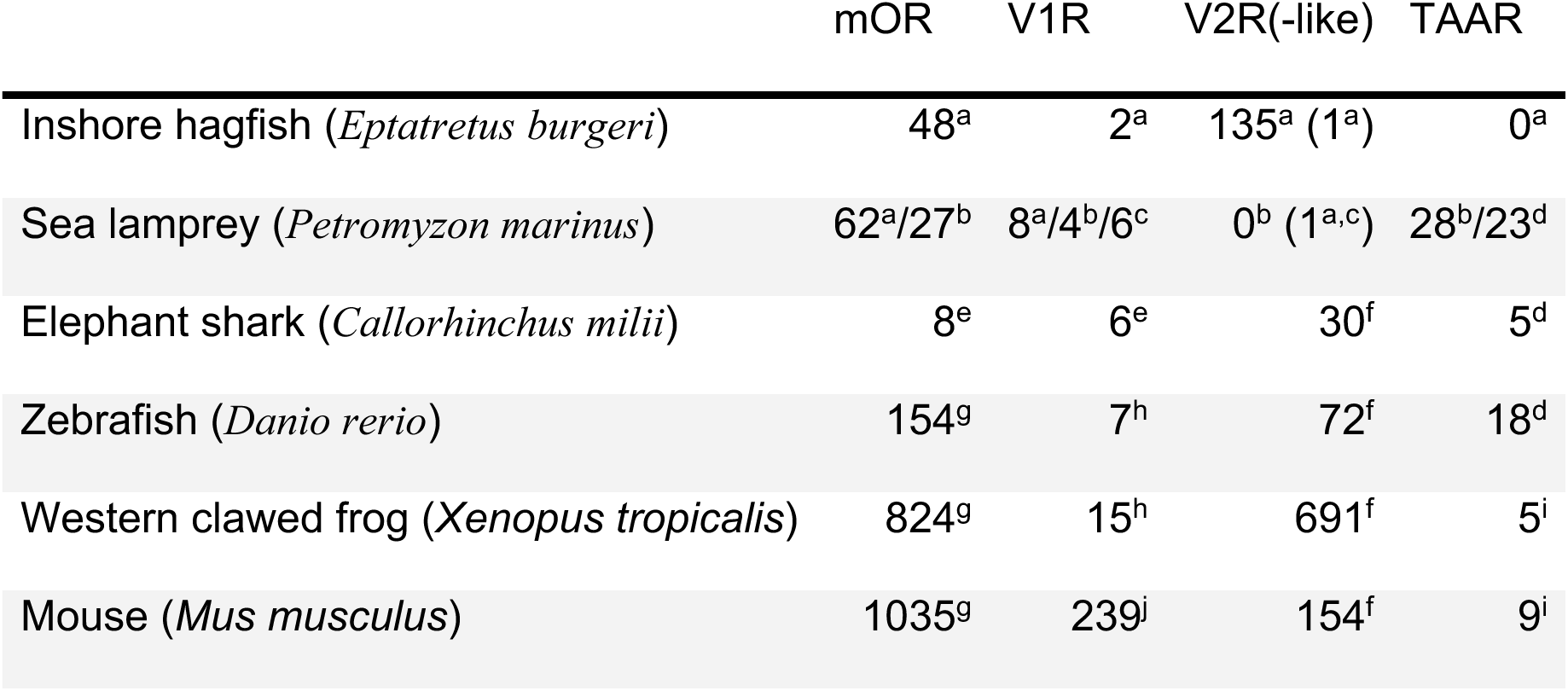
Comparison of OR repertoires in vertebrates.

|  | mOR | V1R | V2R(-like) | TAAR |
| --- | --- | --- | --- | --- |
| Inshore hagfish (*Eptatretus burgeri*) | 48<sup>a</sup> | 2<sup>a</sup> | 135<sup>a</sup> (1<sup>a</sup>) | 0<sup>a</sup> |
| Sea lamprey (*Petromyzon marinus*) | 62<sup>a</sup>/27<sup>b</sup> | 8<sup>a</sup>/4<sup>b</sup>/6<sup>c</sup> | 0<sup>b</sup> (1<sup>a,c</sup>) | 28<sup>b</sup>/23<sup>d</sup> |
| Elephant shark (*Callorhynchus milii*) | 8<sup>e</sup> | 6<sup>e</sup> | 30<sup>f</sup> | 5<sup>d</sup> |
| Zebrafish (*Danio rerio*) | 154<sup>g</sup> | 7<sup>h</sup> | 72<sup>f</sup> | 18<sup>d</sup> |
| Western clawed frog (*Xenopus tropicalis*) | 824<sup>g</sup> | 15<sup>h</sup> | 691<sup>f</sup> | 5<sup>i</sup> |
| Mouse (*Mus musculus*) | 1035<sup>g</sup> | 239<sup>j</sup> | 154<sup>f</sup> | 9<sup>i</sup> |

*Source:* (a) this study; (b) Libants et al. (2009); (c) Kowatschew & Korsching (2022); (d) Dieris et al. (2021); (e) Sharma et al. (2019); (f) Zhang et al. (2022); (g) Niimura (2009); (h) Saraiva & Korsching (2007); (i) Hussain et al. (2009); (j) Young et al. (2010).

#### V1Rs

V1R or olfactory receptor class A (ORA) genes are known to be diversified in sarcopterygians, including tetrapods, in which these receptors are thought to be involved in pheromone detection (Nikaido 2019). We found two V1R candidates in the hagfish genome, while more V1Rs (eight genes) were identified in the sea lamprey genome. It is suggested that vertebrate V1Rs are divided into six subfamilies (ORA1–6; Saraiva and Korsching 2007). Through phylogenetic analysis, we found that the hagfish V1R1 candidate forms a monophyletic clade with gnathostome ORA3/4 along with a sea lamprey V1R, supported by a high bootstrap value (99%; Fig. 1B). The hagfish V1R2 candidate was placed as the sister group of the ORA5 family of gnathostomes with a high bootstrap value (97%). In addition, some sea lamprey V1Rs constructed a clade with gnathostome ORA1/2 with a relatively high support value, although the phylogenetic positions of other sea lamprey genes were unclear due to their relatively low support values.

#### V2Rs

V2Rs are chemoreceptors targeting amino acids and peptide pheromones in gnathostomes (Leinders-Zufall et al. 2004; Kimoto et al. 2005; Speca 1999; Koide et al., 2009; Sato et al., 2005). We identified 137 V2R candidates in hagfish. In contrast, only one V2R-like gene was found in the sea lamprey genome, consistent with previous reports (Kowatschew and Korsching 2022). Phylogenetic analysis revealed that one hagfish V2R candidate forms a clade with lamprey and gnathostome calcium-sensing receptor (CaSR) genes, another grouped with the lamprey V2R-like genes, and the remaining 135 genes forms a monophyletic clade together with high bootstrap value (96%), being situated in a basal position to gnathostome V2Rs more closely than the cyclostome V2R-like clade (Fig. 1C). Based on these results, we estimated the first and second candidate gene as hagfish CaSR and V2R-like, respectively, and the last group containing135 candidates as typical V2Rs.

#### TAARs

Previous studies have reported that TAARs are diversified in lamprey (Libants et al. 2009; Dieris et al. 2021). To find hagfish TAARs, we first performed a reciprocal BLAST search against hagfish gene models, but no plausible candidates were found. For further confirmation, we performed phylogenetic analysis of top hit hagfish sequences and known TAARs. Consequently, these hagfish genes did not show affinity to TAARs but to various groups of GPCRs, including the 5-hydroxytryptamine receptor 4 (5-HT4 or HTR4), adrenoceptor beta 1, and dopamine receptor family (Supplementary File 7). These results suggest that hagfish have no TAAR homologs, consistent with previous research (Guo et al., 2022).

### Genomic distribution of the hagfish ORs

To clarify the diversification pattern of hagfish ORs, we then examined the genomic distribution of the identified genes using the following analyses.

First, we defined phylogenetically related clades for mORs and V2Rs (See Materials and Methods). We categorized three major clades for mORs; Group A corresponded entirely to Clade 1, while Group B was subdivided into Clade 2 and 3 (see Supplementary Fig. 1). For V2Rs, nine major clades (Clade 1–9) were distinguished, and the remaining unclassified genes were categorized into “Else” (see Supplementary Fig. 2).

Next, we mapped all identified OR genes onto the genome (Fig. 2A). As the current *E. burgeri* genome is not assembled at the telomere-to-telomere level, we arranged scaffolds (> 0.1 Mbp) from longer to shorter. As an overall pattern, the OR genes exhibited a patchy distribution. For example, mORs in the same clade tended to co-occur on the same scaffold. Even in each scaffold, phylogenetically related genes tended to be neighboring closely in the genome as follows. We calculated phylogenetic and genomic distances between each pairwise combination of the identified genes, and then made a scatter plot between these two distances (Fig. 2C). As a result, we found that a higher portion of intraclade pairs (32.2%; 140/435 pairs) were located within a shorter genomic distance (<10 Mbp), compared to the interclade pairs (1.0%; 7/693 pairs).

**Figure 2.**
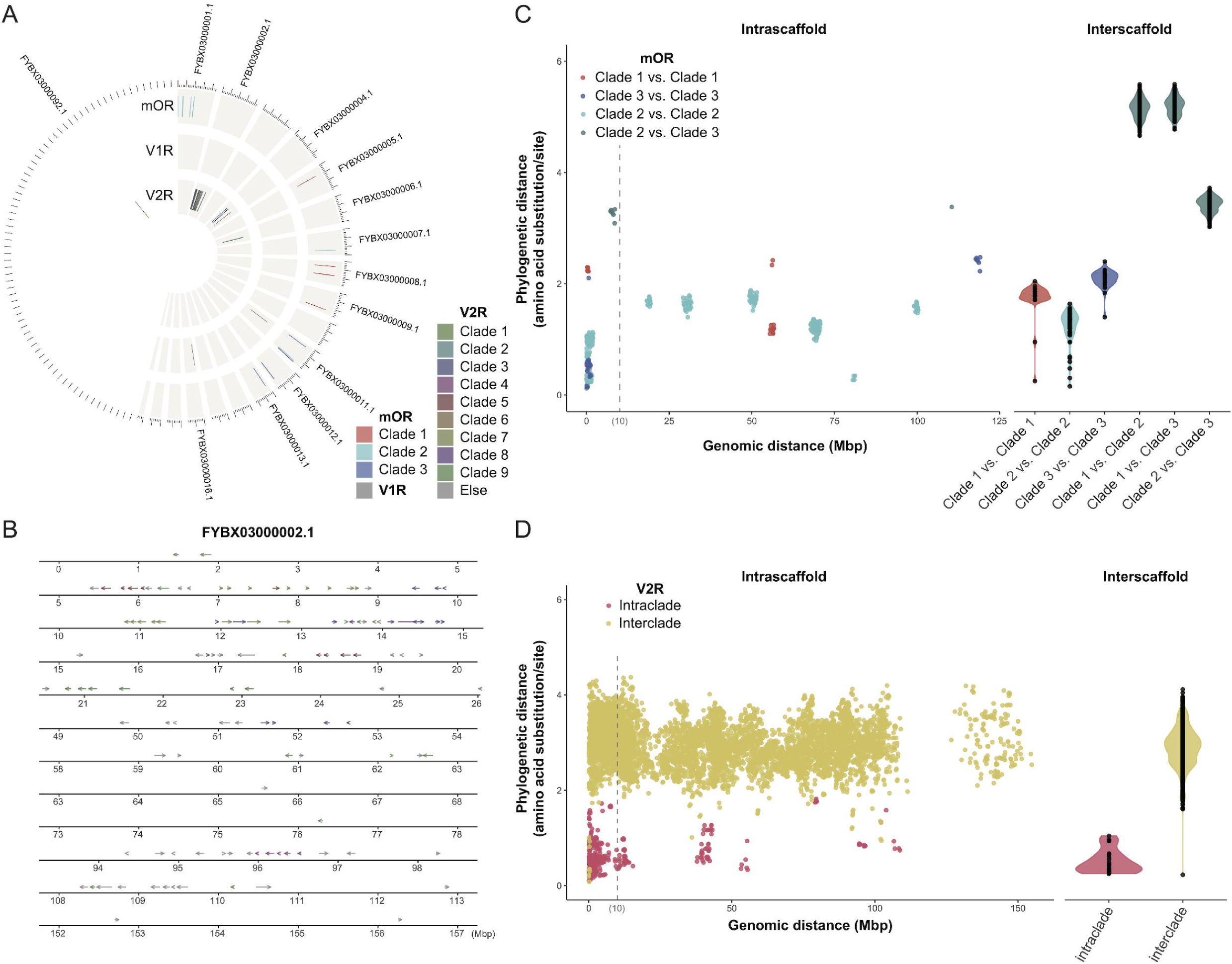
Genomic distribution of the hagfish OR genes. (A, B) Genomic mapping of the OR genes to the scaffolds larger than 0.1 Mbp (A) and to a single scaffold, FYBX03000002.1 (B). Each clade of mOR and V2R genes (for identification of clades, see text) is differently colored. Scale interval: 10 Mbp for (A). (C, D) Scatter plots between phylogenetic and genomic distances, which are calculated between each pairwise combination of the identified genes: mORs (C) and V2Rs (D).

V2Rs also exhibited a patchy genomic distribution, and most of them (91.1%; 123/135 were concentrated on a single scaffold (FYBX03000002.1). In this scaffold, phylogenetically close genes were arranged in clusters along the genome (Fig. 2B). In addition, scatter plotting between phylogenetic and genomic distances (Fig. 2D) showed that most of the intraclade pairs (69.8%; 273/391 pairs) were located within a shorter genomic distance (<10 Mbp), but fewer interclade pairs (21.5%; 1862/8652 pairs) were within such a close distance.

### Organ-specific gene expression of the hagfish ORs

For the examination of whether the hagfish olfactory receptor candidates are indeed expressed in the olfactory organ, we performed organ-specific gene expression analyses using bulk RNA-seq data (the olfactory organ, brain, pituitary gland, gill, intestine, kidney, liver, testis, and ovary). As a result, we confirmed that most of these candidate genes were highly expressed in the olfactory organ.

First, 47 genes among 48 mOR candidates showed high expression levels in the olfactory organ (44 genes showed the highest expression levels in the olfactory organ among the nine tested organs), suggesting their function as true olfactory receptors (Fig. 3). Notably, many (19 genes) are also found to be expressed at a certain level in the testis and/or ovary.

**Figure 3.**
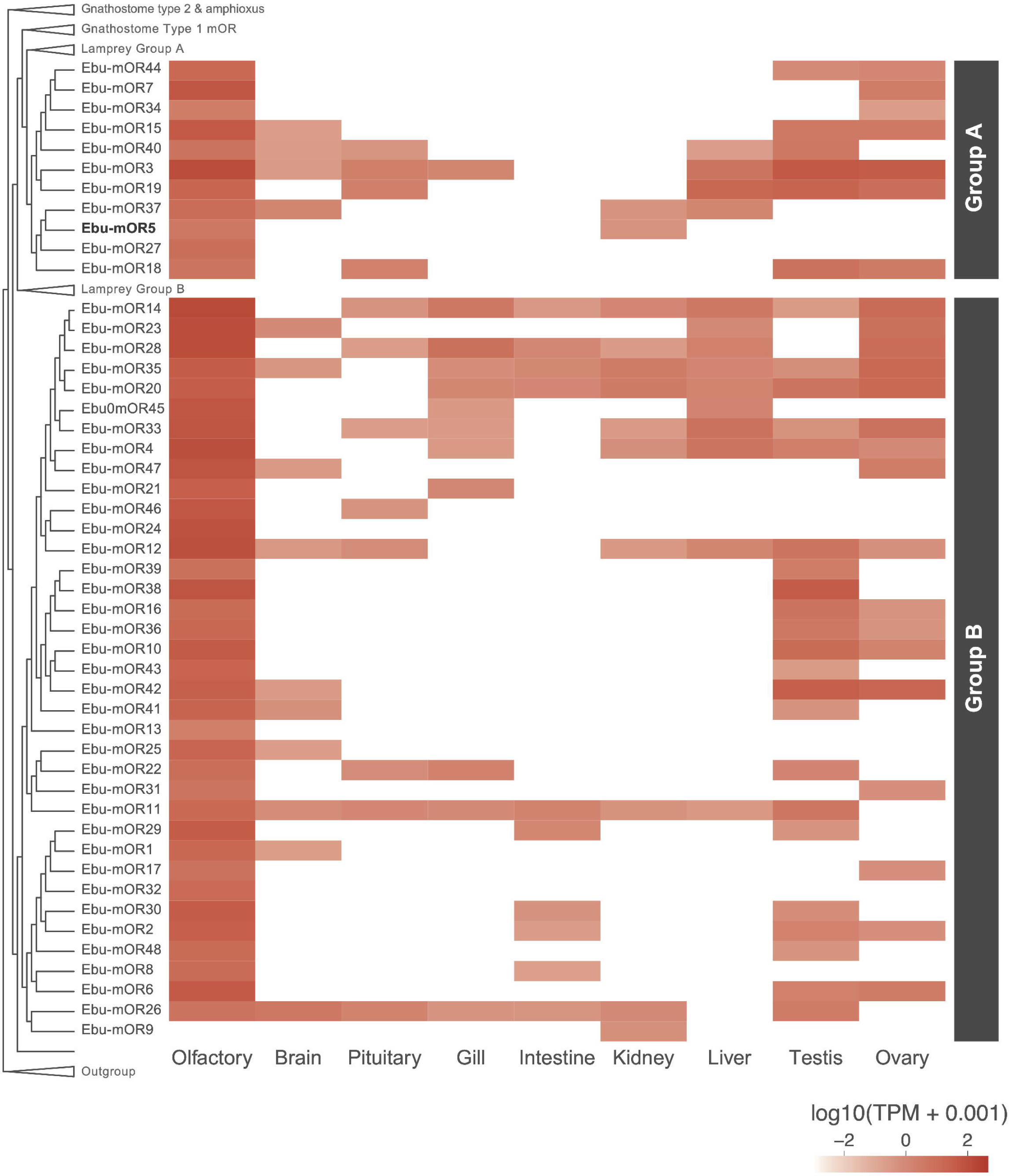
Gene expression profile of mOR candidates across various organs. Expression levels are shown as TPM values, log-transformed. Genes on the y-axis are ordered according to the phylogenetic tree shown in Fig. 1. The mOR gene used for *in situ* hybridization analysis (hagfish mOR5, abbreviated as Ebu-mOR5) is boldfaced.

Second, we confirmed that the two V1R candidates were also expressed at a relatively high level (TPM > 1) in the olfactory organ (Fig. 4A). The Ebu-V1R1 gene was also expressed in the brain at the same expression level, and was slightly weakly expressed in the liver and kidney. The Ebu-V1R2 gene showed the highest expression in the olfactory organ, with lower expression levels in other organs.

**Figure 4.**
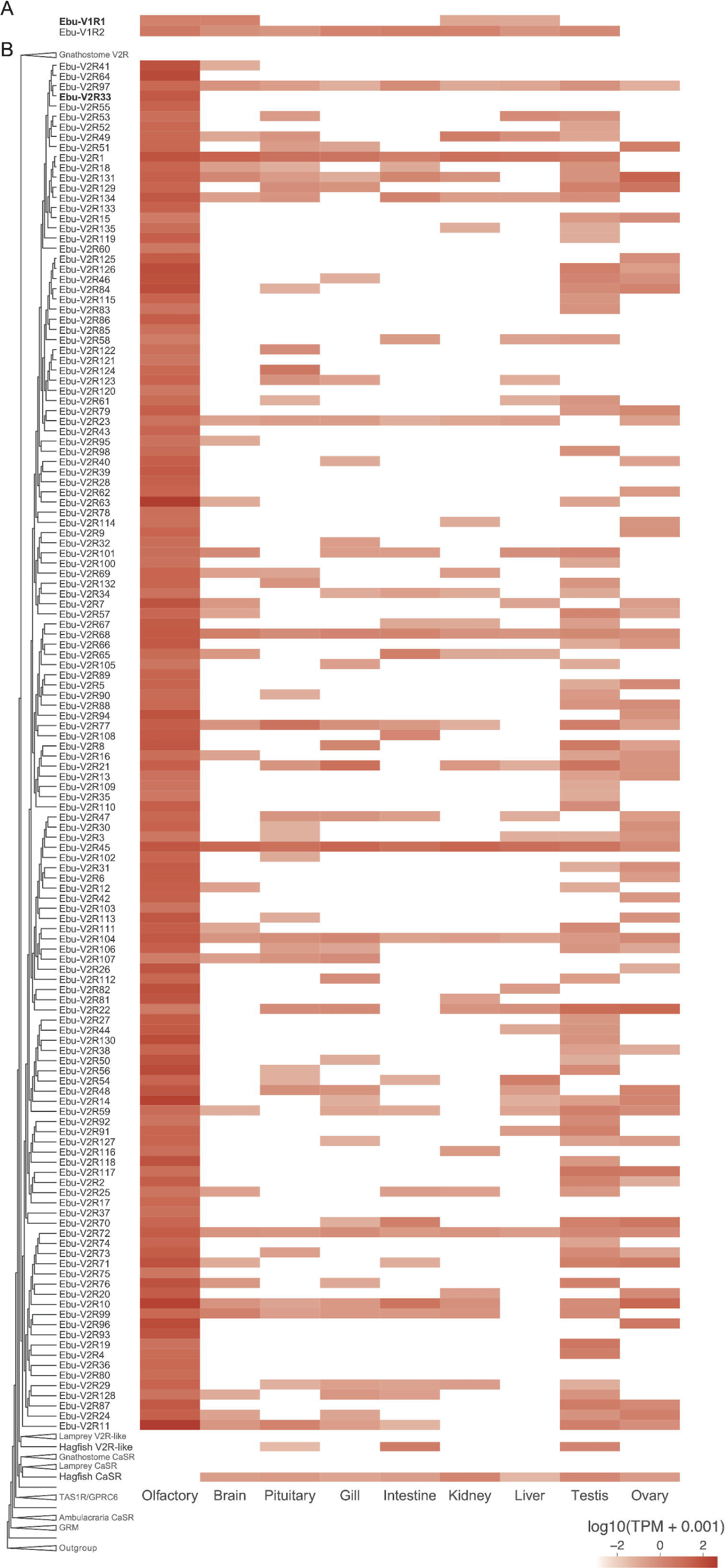
Gene expression profile of organ-specific differential expression analysis for V1R and V2R candidates across various organs. Expression levels are shown as TPM values, log-transformed. Genes on the y-axis are ordered according to the phylogenetic tree shown in Fig. 1. The genes used for *in situ* hybridization analysis (hagfish V1R1 and V2R33, abbreviated as Ebu-V1R1 and Ebu-V2R33, respectively) are boldfaced.

Last, we detected expression of all 135 hagfish putative V2R genes in the olfactory organ (Fig. 4B). Moreover, almost all of them (134 genes except V2R22) showed the highest expression in this organ. Impressively, expression of these genes was also frequently detected in the testis and/or ovary. In contrast, neither the hagfish CaSR nor V2R-like was expressed in the olfactory organ; instead, the former was expressed in all the other organs tested (especially in the kidney and testis), while expression of the latter was detected highly in the intestine and testis, and weakly in the pituitary gland.

### Gene expression patterns of the hagfish ORs in the olfactory epithelium

To investigate whether the olfactory receptor candidates are specifically expressed in olfactory receptor cells, we performed *in situ* hybridization analysis for hagfish mOR (mOR5), V1R (V1R1), and V2R (V2R33), selected based on their expression levels and predicted sequence lengths.

The hagfish olfactory organ, or the nasal basket, is situated anterior to the brain (Fig. 5A; Marinelli and Strenger 1956; Muramatsu et al., 2024). It consists of a series of seven olfactory lamellae, oriented parallel to the body axis and attached to the dorsal roof of the olfactory cavity (Fig. 5B; Døving 1998).

**Figure 5.**
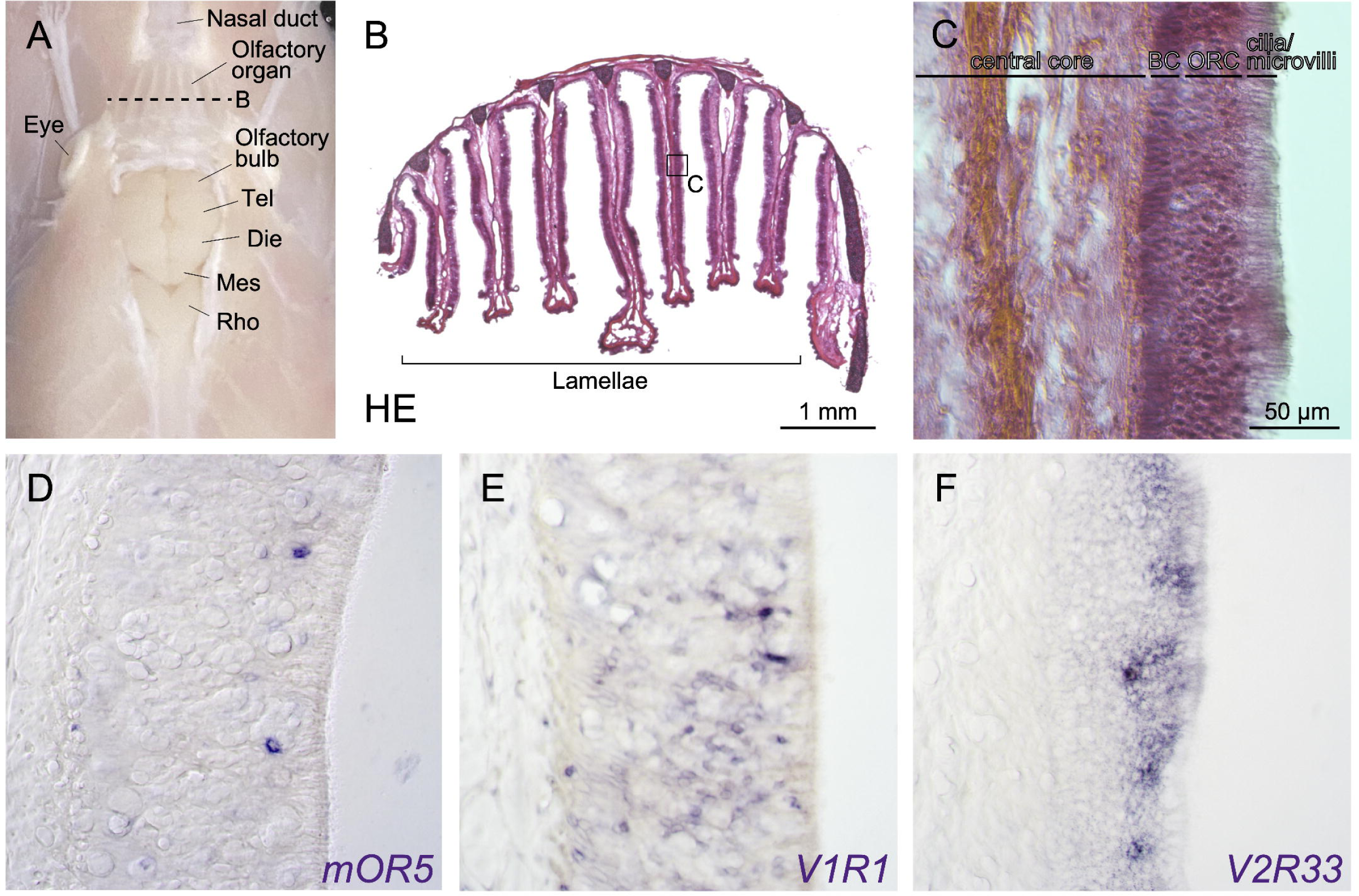
Histological analysis of the hagfish olfactory organ and gene expression patterns of olfactory receptor candidates. (A) Overview of the hagfish olfactory organ and brain, dorsal view. The skin and dorsal roof of the braincase are removed. The section plane for (B) is indicated by the dashed line. (B) Cross-section of the olfactory organ, hematoxylin-eosin (HE) staining. (C) Magnified image of (B), showing the layered structure of the olfactory lamina. (D–E) Gene expression patterns of hagfish *mOR5* (D), *V1R1* (E), and *V2R33* (F). Abbreviations: BC, basal cells; Die, diencephalon; Mes, mesencephalon; ORC, olfactory receptor cell; Rho, rhombencephalon; Tel, telencephalon. The scale bar for (C–F) is shown in (C).

By histological analysis, we first confirmed that the hagfish olfactory lamella exhibits a laminar structure (Fig. 5C), in a manner similar to that of the teleost fish (Hara 1975). Each lamella consists of a medial fibrous layer called the central core and the peripheral olfactory epithelium, which is further subdivided into three layers: from proximal to distal, the basal cell (BC) layer, the olfactory receptor cell (ORC) layer, and the apical cilia/microvilli layer.

Based on this observation, we then examined gene expression patterns of hagfish *mOR5*, *V1R1*, and *V2R33*. As a result, we found that all of them were expressed in olfactory cells in a scattered pattern (Fig. 5D–F). Negative controls using sense probes showed no such expression patterns (Supplementary Fig. 3). Among these genes, V1R1-positive cells showed a distinctive pattern, being broadly and relatively densely distributed in the ORC layer (Fig. 5E). In addition, some *V1R1*-positive cells were also observed in the BC layer. In contrast, the olfactory cells expressing the other two (*mOR5* and *V2R33*) were more sparsely distributed and found in a relatively superficial part of the ORC layer (Fig. 5D, F).

## Discussion

### Hagfish OR repertoire

In this study, we have investigated the olfactory receptor repertoire in hagfish. By combining genome-wide and gene expression analyses, we identified putatively functional olfactory receptor genes as follows (see also Table 1 and Fig. 6).

**Figure 6.**
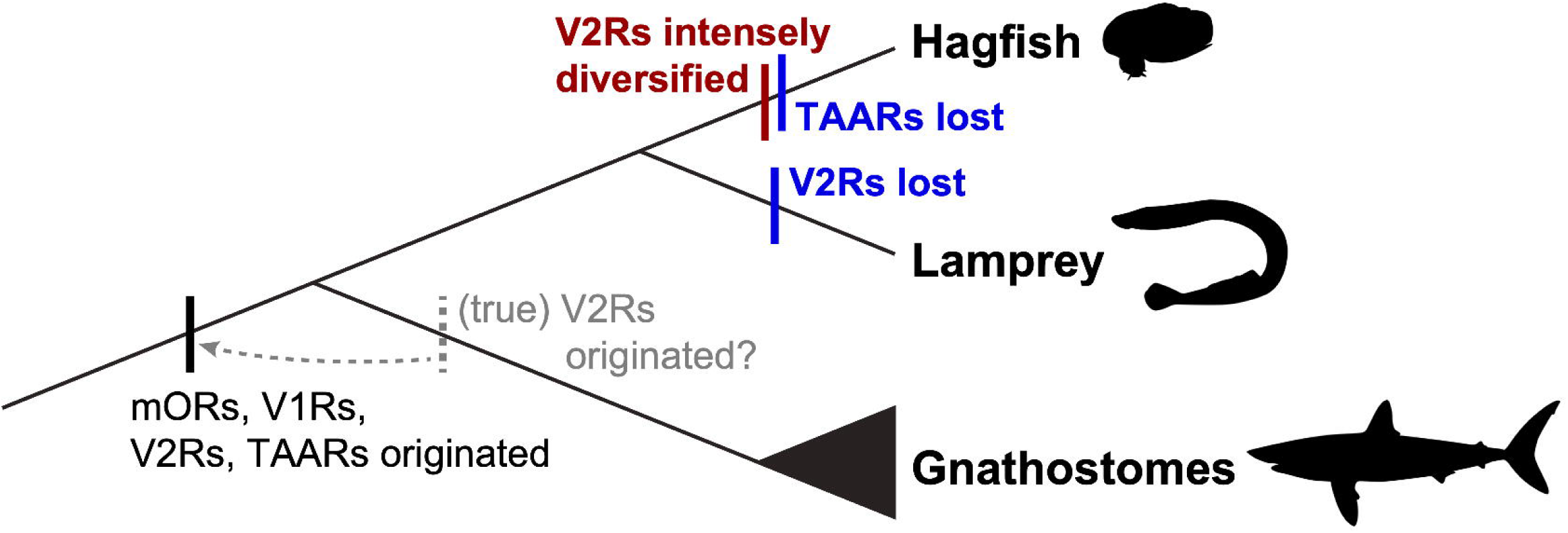
Diversification of ORs in vertebrate evolution. Silhouettes from Phylopic (public domain).

#### mORs

We identified 48 hagfish mOR genes, all of which fall into type 1 mORs (Fig. 1A). In addition, almost all of them (47 genes) are highly expressed in the olfactory organ (Fig. 2), and at least one of them (*mOR5*) in olfactory receptor cells (Fig. 5D), suggesting that these genes encode functional (i.e., “true”) mORs. As these hagfish mORs form a monophyletic group together, these genes are considered to have duplicated through hagfish-specific diversification.

Nevertheless, no hagfish mOR genes were classified as type 2 subfamily. In contrast, the lamprey has both type 1 and type 2 subfamilies (this study; Niimura 2009), suggesting that the common ancestor of all vertebrates had both types, but hagfish has lost type 2 mORs.

#### V1Rs

In this study, we found only two hagfish V1R genes, which show high affinity to the ORA3/4 subfamily and ORA5 subfamily, respectively (Fig. 1B). We also demonstrated that these genes are expressed in the olfactory organ, and at least one of them (*V1R1*) is expressed particularly in olfactory receptor cells (Fig. 4A, 5E). These results suggest that hagfish have true V1Rs.

Still, we identified more V1Rs (eight genes) from the latest version (Timoshevskaya et al., 2023) of the sea lamprey genome. These genes include two additional ones that were unreported in previous research (Kowatschew and Korsching 2022). Among them, two (lamprey V1R1 and V1R2) show affinity to the gnathostome ORA1/2, two (lamprey V1R3 and V1R7) to the gnathostome ORA3/4, and one (lamprey V1R6) to ORA6. The remaining three (lamprey V1R4, V1R5, and V1R8) were placed in the basal position of the gnathostome ORA5/6, despite showing low support values. Combining these results with previous research (Kowatschew and Korsching 2022), it is suggested that the common ancestor of all vertebrates had at least three subtypes of V1Rs, ORA1/2, ORA3/4, and ORA5/6, and that the hagfish has lost the ORA1/2 gene(s).

#### V2Rs

Most notably, we revealed that V2Rs show the highest diversification among hagfish olfactory receptors. We identified typical V2Rs (135 genes) that form a monophyletic clade basal to the gnathostome V2Rs. As the closest outgroup to these V2Rs, we found a pair of V2R-like genes in hagfish and lamprey. Consistent with a previous suggestion that lamprey V2Rs do not function as olfactory receptors (Kowatschew and Korsching 2022), our transcriptome analysis indicates that the hagfish V2R-like gene is not expressed in the olfactory organ at a functional level (Fig. 3B).

In contrast, we confirmed that all of the true V2Rs are expressed in the olfactory organ (Fig. 3B), particularly and at least one of them (*V2R33*) in olfactory receptor cells (Fig. 5F), suggesting that these genes do serve as olfactory receptors. This finding challenges the conventional view that the olfactory function of the V2Rs was established in the gnathostome lineage (Kowatschew and Korsching 2022; Zhang et al. 2022). Instead, we suggest that true V2Rs had already been acquired in the common ancestor of all vertebrates; these genes underwent extensive diversification in hagfish, but they have been lost specifically in the lamprey lineage.

#### TAARs

We found no TAAR gene candidates in the current version of the hagfish genome, consistent with previous research (Guo et al., 2022). In contrast, previous studies (Libants et al. 2009; Dieris et al. 2021) reported that the lamprey has dozens of TAARs. Therefore, it is suggested that TAARs can be traced back to the common ancestor of all vertebrates and that they have been lost specifically in the hagfish lineage.

### Importance of assembly quality for comparative genomics

To obtain a more complete gene list for comparison, we re-surveyed lamprey ORs using the latest version (Timoshevskaya et al., 2023) of the sea lamprey genome. As a result, we identified a greater number of mOR and V1R genes than previously reported (Table 1), suggesting that better genome assemblies provide more reliable information for comparative genomics.

As the current version of the *E. burgeri* genome used in this study is not completely assembled at the telomere-to-telomere level (Yu et al., 2024), future high-quality assemblies may reveal additional OR genes that are not detected in this study. In addition, if a complete genome assembly were available, our analysis of the genomic distribution of hagfish OR genes would be more precise. Still, the overall trends in OR gene diversification, loss, and genomic distribution are clearly demonstrated.

### Gene duplication of hagfish ORs

Our analysis of the genomic mapping of hagfish OR genes revealed that phylogenetically close mOR and V2R genes tend to reside close in the hagfish genome. Particularly in the scaffold FYBX03000002.1, phylogenetically close V2R genes were organized in clusters along the genome. However, a given genomic cluster often contains genes from different clades (Fig. 2B). This distribution pattern suggests that these genes have been diversified through tandem gene duplications and chromosomal rearrangements, as has been observed in human OR gene evolution (Niimura and Nei 2005).

It is suggested that whole-genome duplication (WGD) may contribute to the genetic diversity of the ORs (Wang et al. 2021). Recently, the cyclostomes were proposed to have experienced lineage-specific WGD or triplication (Marlétaz et al., 2024; Yu et al., 2024).

Nevertheless, this event appears to have had no or a small contribution to the evolution of ORs, especially V2Rs. In fact, the scaffold FYBX03000002.1 includes most of the hagfish V2R genes (91%), indicating that these genes have duplicated at the intrachromosomal level.

Another possible mechanism is retrotransposon-mediated duplication. However, this scenario is unlikely in the diversification of hagfish V2Rs because retrotransposon-encoded reverse transcriptase typically generates intronless genes (Lallemand 2020), and all hagfish V2Rs contain introns except for a few exceptions whose domains were only partially predicted (Supplementary File 1).

Taken together, the most plausible mechanism for the extensive diversification of hagfish V2Rs is tandem gene duplications followed by chromosomal rearrangements.

### Expression specificity of the ORs

In mammals, it is widely believed that each olfactory receptor neuron expresses only one OR gene (Bystrova & Kolesnikov, 2021). This “one neuron–one receptor” hypothesis was originally proposed based on studies in mice and rats (e.g., Chess et al., 1994; Malnic et al., 1999). Nevertheless, it has yet to be conclusively demonstrated (Mombaerts, 2004). In fact, a previous study showed that two functional odor receptors are expressed in one neuron in the fruit fly (Goldman et al., 2005).

Particularly, whether the “one neuron–one receptor” rule also applies to basal vertebrates remains an open question. The broad expression pattern of the V1R gene in the hagfish olfactory epithelium (Fig. 5E) imply the possibility that V1Rs are coexpressed with other ORs in a single receptor neuron. To test this hypothesis in cyclostomes, further analyses—such as single-cell RNA-seq—are necessary.

### Cell morphology and distribution of the OR-expressing neurons

Certain vertebrates exhibit lineage-specific distribution patterns of olfactory receptor cells. For example, mammals have two distinctive olfactory organs: the main olfactory organ and the vomeronasal organ. The former consists of mOR- and TAAR-expressing ciliated cells, while the latter is specialized for pheromone detection and contains V1R- and V2R-expressing—and additionally in rodents (Boillat et al., 2021), formyl peptide receptor (FPR)-expressing—microvillous cells (Poncelet and Shimeld, 2020). In teleost fish, there are four types of olfactory receptor cells: mOR- and TAAR-expressing ciliated cells, V1R- and V2R-expressing microvillous cells, V1R-expressing crypt neurons, and kappe neurons with unidentified receptors (Poncelet & Shimeld, 2020). Among them, the mOR-expressing ciliated cells are found in the deep layer of the olfactory epithelium, while the V1R/V2R-expressing microvillous cells, along with crypt neurons, are located in the superficial layer (Hansen et al., 2004; Sato et al., 2005).

In hagfish, ciliated and microvillous cell types have been reported to be present (Døving, 1998). However, the relationship between these cell types and OR expression patterns, as well as the spatial distribution of olfactory receptor cells, remain unresolved. Although our gene expression analysis suggests that both mOR- and V2R-expressing cells are located in the relatively superficial layer (Fig. 5D, F), further detailed analyses, including spatial RNA-seq, would provide key information for these questions.

### Lineage-specific diversification of olfaction in early vertebrates

One of the most notable findings of this study is that functional V2Rs are present and extensively diversified in the cyclostome hagfish. While V2Rs are reported to bind to water-soluble molecules such as the peptide ligands of MHC class I molecules and exocrine gland peptides (ESPs) in mammals (Leinders-Zufall et al. 2004; Kimoto et al. 2005), they are also expected to detect amino acids and their derivatives, eliciting feeding behaviors in teleosts (Speca 1999; Koide et al., 2009; Sato et al., 2005). The massive diversification of hagfish V2Rs may thus reflect an adaptation to scavenging life in low-light environments, with various functions such as the detection of food sources and mating partners. To determine the functions of hagfish V2Rs, it is necessary to examine these receptors from multiple disciplinary perspectives, including ligand binding assays, electrophysiological recordings, and behavioral experiments using candidate odorants.

Furthermore, our phylogenetic analyses revealed that the OR repertoires in basal vertebrates have evolved in a mosaic manner (Fig. 6). On the one hand, hagfish have experienced extensive diversification of V2Rs and loss of TAARs. On the other hand, lampreys appear to have lost true V2Rs. As previous studies have relied on lampreys only, it has been mistakenly suggested that true V2Rs originated in the common ancestor of gnathostomes. However, this study pushes the evolutionary origin of the V2Rs back to the common ancestor of all vertebrates. This finding calls attention to the importance of hagfish, which has often been overlooked, yet holds significant potential to illuminate critical aspects of early vertebrate evolution.

## Conclusion

In this study, we conducted a comprehensive genome-wide survey and gene expression analyses of ORs in hagfish, a basal vertebrate that typically inhabits the deep sea. We identified 48 mORs, two V1Rs, 135 typical V2Rs, and no TAARs in the inshore hagfish (*E. burgeri*). Phylogenetic analysis revealed that all mORs belong to the type 1 subfamily. The two V1Rs form a clade with known vertebrate ORA subfamilies, and the 135 V2Rs form a distinct monophyletic clade basal to gnathostome V2Rs. Genomic mapping revealed large clusters of the hagfish V2Rs on a single scaffold, likely resulting from tandem gene duplications and chromosome rearrangements. Gene expression profiling confirmed that nearly all of these receptors are specifically expressed in the olfactory organ, and *in situ* hybridization showed that at least some are localized in olfactory receptor cells.

These findings revise the current evolutionary scenario of vertebrate olfactory systems. While lampreys lack typical V2Rs, our data indicate that these receptors were already present in the common ancestor of vertebrates and were subsequently lost in the lamprey lineage. The large-scale expansion of V2Rs in hagfish likely reflects ecological adaptation to their scavenging niche in dark environments. Our results elucidate the previously unrecognized evolutionary diversity in the olfactory system of basal vertebrates, highlighting hagfish as a key animal for understanding the early vertebrate evolution.

## Acknowledgment

We thank Drs. Hiroshi Wada and Yoshiaki Morino for their valuable comments, and Dr. Haruka Ozaki for providing computer resources. We appreciate Susumu Hatanaka (Shinsho Maru; Fujisawa, Kanagawa, Japan) and Masanori Nishizaki and Dr. Masa-aki Yoshida (Oki Marine Biological Station, Shimane University) for providing the *E. burgeri* specimens.

Computations were partially performed on the NIG supercomputer at ROIS National Institute of Genetics. This work was supported by the Grant-in-Aid for the Japan Society for the Promotion of Science (JSPS; Grant Numbers JP20K15855, JP22K15164, JP24K09556, and JP24H01538 to D.G.S, and JP19K16178 to Y.Y.) and by the Sasakawa Scientific Research Grant from The Japan Science Society (Grant Number 2023-4098 to H.K.).

## Materials and methods

### Animals

Inshore hagfish (*E. burgeri*) specimens, both male and female, were collected from Sagami Bay, Kanagawa Prefecture in 2021–2025 or from Kamo Bay near Oki Marine Biological Station, Shimane University (Oki, Shimane, Japan) in 2021. As for the animals from Sagami Bay, they were transported to the laboratory at the University of Tsukuba, Japan, where they were kept in 160 L tanks filled with circulating artificial seawater at 12°C ± 0.5 °C before dissection. As for the animals from Kamo Bay, they were transported to a laboratory at Matsue campus, Shimane University (Matsue, Shimane, Japan). They were maintained in a 260 L tank filled with circulating artificial seawater at 15 ± 0.5 °C and sacrificed for tissue sampling. All procedures in this study were performed in compliance with the guidelines for animal use of the Animal Care Committees at the University of Tsukuba and Shimane University (specific approval is not required for experimentation on fishes under the Japanese law, Act on Welfare and Management of Animals). During the investigation, every effort was made to minimize suffering and to reduce the number of animals used.

### Tissue sampling

For RNAseq of the olfactory organ, the tissue was collected from the decapitated animal following ice anesthesia, and was frozen using liquid nitrogen and kept at −80 °C until analyzed. For molecular cloning, histological analysis, and *in situ* hybridization, animals were deeply anesthetized with MS-222 (100 mg/L; Sigma, A5040) and euthanized by decapitation. Then, the decapitated heads were dissected to harvest the olfactory organs, which were served for each experiment described below.

### RNAseq and differential gene expression analysis

RNA-seq of the olfactory organ was newly performed in this study, following the method for other organs described previously (Yamaguchi et al., 2023). Total RNAs were extracted from the frozen tissue from a single individual (total length 55 cm, weight 226.3 g, male; as for other tissues, the pituitary glands from five individuals of mixed sexes and sizes were pooled, while other tissues were collected from a single individual, as reported by Yamaguchi et al., 2023). Raw RNA-Seq data are available at the public repository (BioProject accession ID: PRJDB35770, DRR Run: DRR707969–DRR707972, DRR707975-DRR707977, DRR707979, and DRR707980).

The adaptor sequences and low-quality sequences were then removed using Trimmomatic version 0.38 (Bolger et al., 2014). We used FastQC version 0.11.9 (Andrews, 2010) to evaluate the sequence data quality. The gene models of *E*. *burgeri* were retrieved from Ensembl Release 104 and were used as a reference for calculating expression levels. To prepare a more comprehensive reference dataset, we added sequences identified from the genome by TBLASTN searches. The expression amount was quantified as transcripts per kilobase million (TPM) using Salmon version 1.10.3 (Patro et al., 2017). For visualizing the expression amount of olfactory receptor genes, we used the tidyverse and ggplot2 packages in R. TPMs were log10-transformed after adding a pseudocount of 0.001.

### Gene search

To identify main olfactory receptor (mOR) gene candidates, we performed BLASTP searches on the hagfish and sea lamprey (*P. marinus*) gene models using mOR sequences of gnathostomes as queries. The gnathostomes included the following species: human, mouse, zebrafish, and African clawed frog. We retrieved hagfish gene models and genome sequences from ENSEMBL (Release 104). As a new version of the sea lamprey genome was reported recently (Timoshevskaya et al., 2023), we re-surveyed mORs of this animal. The amino acid (AA) sequences of the gnathostomes were retrieved from Niimura and Nei (2005). We extracted hagfish and sea lamprey sequences from the BLAST search result with E values < 1E-3 and an alignment length of ≥ 250 AA. To confirm the homology, we also conducted BLASTP searches against the human, zebrafish, and African clawed frog gene models (ENSEMBL release 104) using the hagfish and sea lamprey sequences as queries with the same thresholds. For gene extraction, we used SeqKit ver. 2.8.2 (Shen et al., 2024). To narrow down the list to more plausible sequences, we performed a domain search using HMMER 3 (ver. 3.2.1) with the Pfam database (ver. 34.0). As a result, we found 35 genes from the hagfish and 61 genes from the sea lamprey gene models.

Recently, a new version of the *E. burgeri* genome was published (Yu et al., 2024), but gene models based on this genome data are not officially provided. To extract potential mOR gene candidates that were missed in the gene models based on the old version, we searched for mOR candidate genes directly from the new one. To identify potential mOR gene candidates, TBLASTN was performed on the hagfish genome version 4.0 (https://www.ncbi.nlm.nih.gov/datasets/genome/GCA_900186335.3/). We used the same query sequences as the analysis using gene models. The threshold for homologous sequences was E value < 1E-10 and alignment length ≥ 250AA as described in the previous research (Niimura and Nei, 2005). To confirm their homology, we also conducted BLASTP searches as described above. The coding sequences were predicted from TBLASTN hit regions with 900 bp of flanking sequences using GeneWise (version 2.4.1) with default parameters. We used the same query sequences for this TBLASTN search as those used for the BLASTP search described above. Next, the possibly functional sequences were extracted from the predicted sequences using HMMER 3 and the Pfam database. 29 sequences were matched to the mOR candidate genes extracted from the gene models, and 13 sequences were newly identified from the genome sequences.

We collected candidate sequences of hagfish V1R and V2R genes in the same way as for mORs. In the reciprocal BLAST search of gene models, homologous genes of V2R, CaSR, taste receptor type 1 (TAS1R), and G protein-coupled receptor family C group 6 (GPRC6) were extracted as the candidate sequences for V2R. The plausible sequences were extracted with at least 200 AA of a 7-transmembrane domain (PF00003, TM7_3) by domain searches for V2R. As a result of BLAST searches and domain searches against the gene models, two and 38 gene models were identified as candidates of V1R and V2R, respectively. The one and 99 additional candidate sequences of V1R and V2R were found from the hagfish genome in a similar way to identify mOR sequences. To predict coding sequences by GeneWise, we used sequences found by TBLASTN against the TM7 domain. As V2Rs contain another domain in their N-terminal region, we also included this domain and the intron sequences for prediction if a TBLASTN hit sequence is found to contain it (on the same strand and intersequences between the TBLASTN hits < 100,000 bp). In the other case, that is, even if a TBLASTN hit sequence does not include the N-terminal region, we also used any such sequence for our analysis because there is a possibility that its N-terminal domain is just not detected. In addition, we identified 11 more plausible sequences that correspond to the gene models. Thus, we used 110 predicted genes, including 11 sequences for downstream analysis.

For lamprey V1R genes, we initially identified four genes from the sea lamprey (*P. marinus*) gene models. While three genes were previously reported (Kowatschew and Korsching, 2022), the other one (Pma-V1R7) was newly identified. Because the detected genes were fewer than V1Rs reported in the previous study, we further conducted a TBLASTN search against the sea lamprey genome. As a result, we detected seven V1R candidate sequences (six known V1R genes and Pma-V1R7). Additionally, we searched V1R sequences against the arctic lamprey (*L. camtschaticum*) genome and gene models. We detected the seven genes, of which five correspond to the genes identified in the previous research. Although another gene corresponded to Pma-V1R7, it was truncated. One of the seven genes (Lca-V1R8) was newly identified, and we also found the orthologous gene (Pma-V1R8) from the sea lamprey genome. In summary, we identified eight lamprey V1R genes for the phylogenetic analysis.

### Phylogenetic analysis

We retrieved mOR sequences of the gnathostomes and the amphioxus from Niimura (2009), V1R and TAS2R sequences from the ENSEMBL database and previous research (Kowatschew and Korsching, 2022, Sharma et al., 2019, Zapilko and Korsching, 2016, Date-Ito et al., 2008), and V2R, TAS1R, GPRC6, CaSR, glutamate receptor metabotropic (GRM), and gamma-aminobutyric acid type B receptor (GABBR) sequences from previously published papers and the ENSEMBL database (Zhang et al., 2022, Kowatschew and Korsching, 2022). These sequences were aligned using MAFFT version 7.481 (Katoh and Standley 2013) with the default option and trimmed to poorly aligned regions using TrimAL version 1.4. rev15 (Capella-Gutiérrez, Silla-Martínez, and Gabaldón 2009) with the “gappyout” option. Phylogenetic analysis was conducted using RAxML version 8.2.12 (Stamatakis 2014) with the options of “-f a -x 12345 -p 12345 -# 500 -m PROTGAMMAAUTO --auto-prot=aic -T 16.”

### Genomic mapping and distribution analysis

To classify subgroups of mOR genes and V2R genes, we focused on clades containing at least five genes with their bootstrap value ≥ 85%. We aimed to obtain three or more clades for mORs and V2Rs, respectively, trying to define as larger clades as possible at the same time. Consequently, we obtained three clades for mORs and nine clades for V2Rs (Supplementary Fig.1 and 2). The V2R genes that were not classified into any of these clades were categorized into “Else” (Supplementary 2). We then visualized the genomic distribution of the OR genes using ggbio version 1.46.0 within R version 4.2.2 (R Center For Statistical Computing, Vienna, Austria), based on this classification.

For scatter plotting, we calculated the phylogenetic and genomic distances for all interclade and intraclade pairs of the mOR genes and of the V2R genes, respectively, based on the clade classification. We included the pairs of “Else” V2R genes into the interclade pairs. The phylogenetic distances were obtained using the “cophenetic.phylo” function in the ape version 5.7.1. Accordingly, we visualized the relationship between the genomic and phylogenetic distances using the tidyverse version 2.0.0 and ggplot2 version 3.5.2.

### Molecular cloning and probe synthesis

Total RNA was extracted from the harvested olfactory organs of adult *E. burgeri* using TRIzol Reagent (Invitrogen, 15596026). These RNAs were reverse transcribed into cDNAs using PrimeScript II 1st strand cDNA Synthesis Kit (Takara, 6210A) and were used as templates in the polymerase chain reaction (PCR). Hagfish genes of interest were amplified by PCR using the primers listed in Supplementary Table 1. We used reverse primers including the T3 promoter sequence (20 bp) to synthesize DIG-labeled RNA probes for *in situ* hybridization. As a negative control, we instead used forward primers including the T7 promoter sequence (20 bp) to synthesize DIG-labeled RNA probes for *in situ* hybridization (sequences not shown). DIG-labeled RNA probes were transcribed from PCR products using T3 and T7 RNA polymerase (Roche, 11031163001 and 12352204, respectively).

### Hematoxylin-eosin (HE) staining and section *in situ* hybridization

To perform histological analysis and *in situ* hybridization, the harvested olfactory organs were fixed with 4% PFA/PBS for 1 hour at room temperature (RT) or overnight at 4 C°. The fixed specimen was then rinsed with PBS, dehydrated in a graded methanol series (25%, 50%, 75%, and 100%), and stored at −20 °C.

For cryosectioning, the stored samples were rehydrated in a graded methanol series (75%, 50%, and 25% in PBS) and PBS. They were replaced with 10% and then 30% sucrose. Subsequently, the specimens were embedded in Tissue-Tek Optimum Cutting Temperature (O.C.T.) Compound (Sakura Finetek Japan, 4583). Frozen sections in 20 μm were prepared using a cryostat (Leica, CM1860) and mounted on MAS-coated slide glasses (Matsunami Glass Ind., SMAS-01).

HE staining was performed according to the conventional method after sections were washed with PBS to remove the O.C.T. compound. Namely, the sections were incubated in Mayer’s hematoxylin solution (Wako 131-09665) for 15 min and then rinsed with running tap water for 30 min. Afterward, the sections were incubated in 1% Eosin Y solution (Wako 051-06515). The stained sections were dehydrated in a graded methanol series (25%, 50%, 75%, 95%, and 100%) and xylene-equivalent solvent G-NOX (Genostuff, GN04), and mounted using xylene-free mounting medium PARAmount-D (Falma, 308-500-1).

For *in situ* hybridization, the sections were first washed with PBS to remove the O.C.T. compound and postfixed for 20 min with 4% PFA/PBS and rinsed with PBS. Afterward, they were prehybridized in hybridization buffer (50% Formamide, 5× SSC, 5× Denhardt’s solution, 50 μg/ml yeast tRNA, and 50 μg/ml heparin in deionized-distilled H_2_O) and then incubated in the hybridization buffer with 0.1 mg/ml probe overnight at 60 °C. Subsequently, the specimens were washed with the following solutions: 50% Formamide and 2× SSC (for 30 min at 60 °C, twice), 2× SSC (for 30 min at 60 °C, twice), 0.2× SSC (for 30 min at 60 °C, twice), and PBS (5 min at RT). For signal detection, the specimens were blocked with 0.5% blocking reagent (Roche, 11277073910) in PBS for 1 hour at RT and then incubated with 0.5 μl/ml alkaline phosphatase (AP)-conjugated anti-digoxigenin Fab fragments (diluted in 0.5% blocking reagent/PBS; Roche, 11093274910) overnight at 4 °C. After washing with tris-buffered saline (TBS) four times for 30 min each at RT, alkaline phosphatase signals were detected with 20 μl/ml NBT/BCIP in color development buffer (100 mM Tris HCl pH 9.5, 100 mM NaCl, and 50 mM MgCl_2_). The stained specimens were postfixed in 4% PFA/PBS, dehydrated in a graded methanol series (25%, 50%, 75%, 95%, and 100%) and G-NOX, and mounted using PARAmount-D. The prepared slides were examined under a microscope (Nikon, ECLIPSE Ni) and photographed using a microscope digital camera (Nikon, DS-Ri1).

## Figure Legend

**Supplementary Figure 1.** Classification for mOR genes into Clade 1–3.

**Supplementary Figure 2.** Classification for V2R genes into Clade 1–9.

**Supplementary Figure 3.** Negative controls for *in situ* hybridization using sense probes of *mOR5* (A), *V1R1* (B), and *V2R33* (C). The scale bar for (A–C) is shown in (A).

## Table

**Supplementary Table 1.**
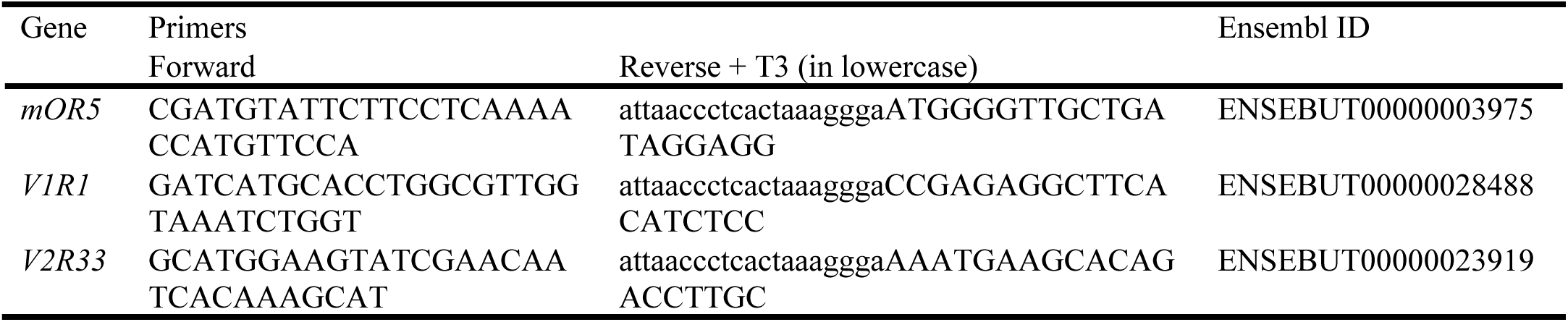
Primer sequences for RNA probes.

## Supplementary Files

**Supplementary File 1.** Details of identified hagfish genes.

**Supplementary File 2.** Details of identified sea lamprey genes.

**Supplementary File 3.** Gene list used for phylogenetic analysis.

**Supplementary File 4.** Detailed phylogenetic tree of mOR genes.

**Supplementary File 5.** Detailed phylogenetic tree of V1R genes.

**Supplementary File 6.** Detailed phylogenetic tree of V2R genes.

**Supplementary File 7.** Phylogenetic tree of known TAAR genes with their close GPCR families, including the 5-hydroxytryptamine receptor 4 (5-HT4 or HTR4), adrenoceptor beta 1, and dopamine receptor genes.

