## Supplementary figures and images for "Genome-wide and gene expression analyses of the hagfish olfactory receptors illuminate lineage-specific diversification of olfaction in basal vertebrates"

### Supplemental Fig. 1

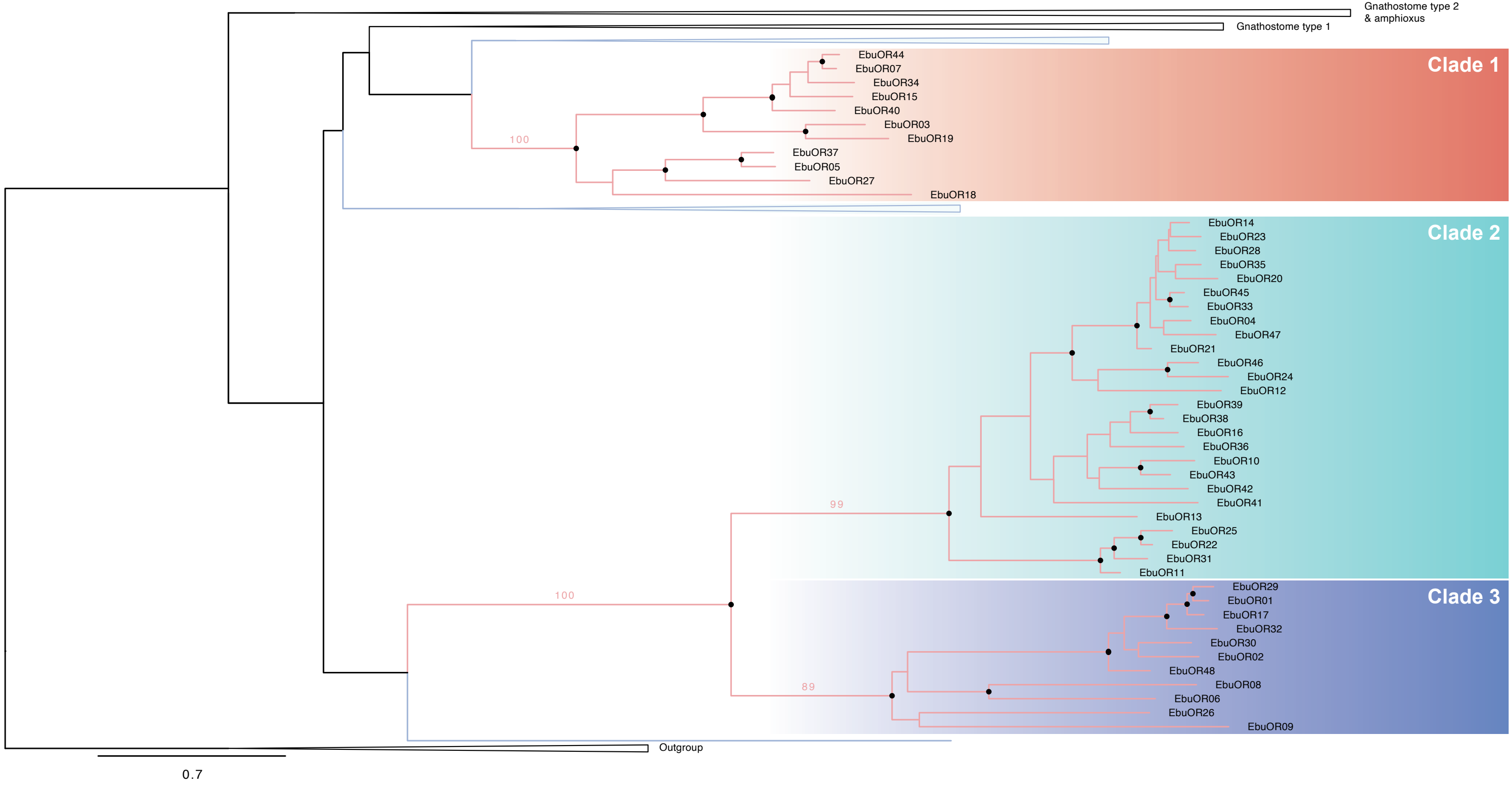

### Supplemental Fig. 2

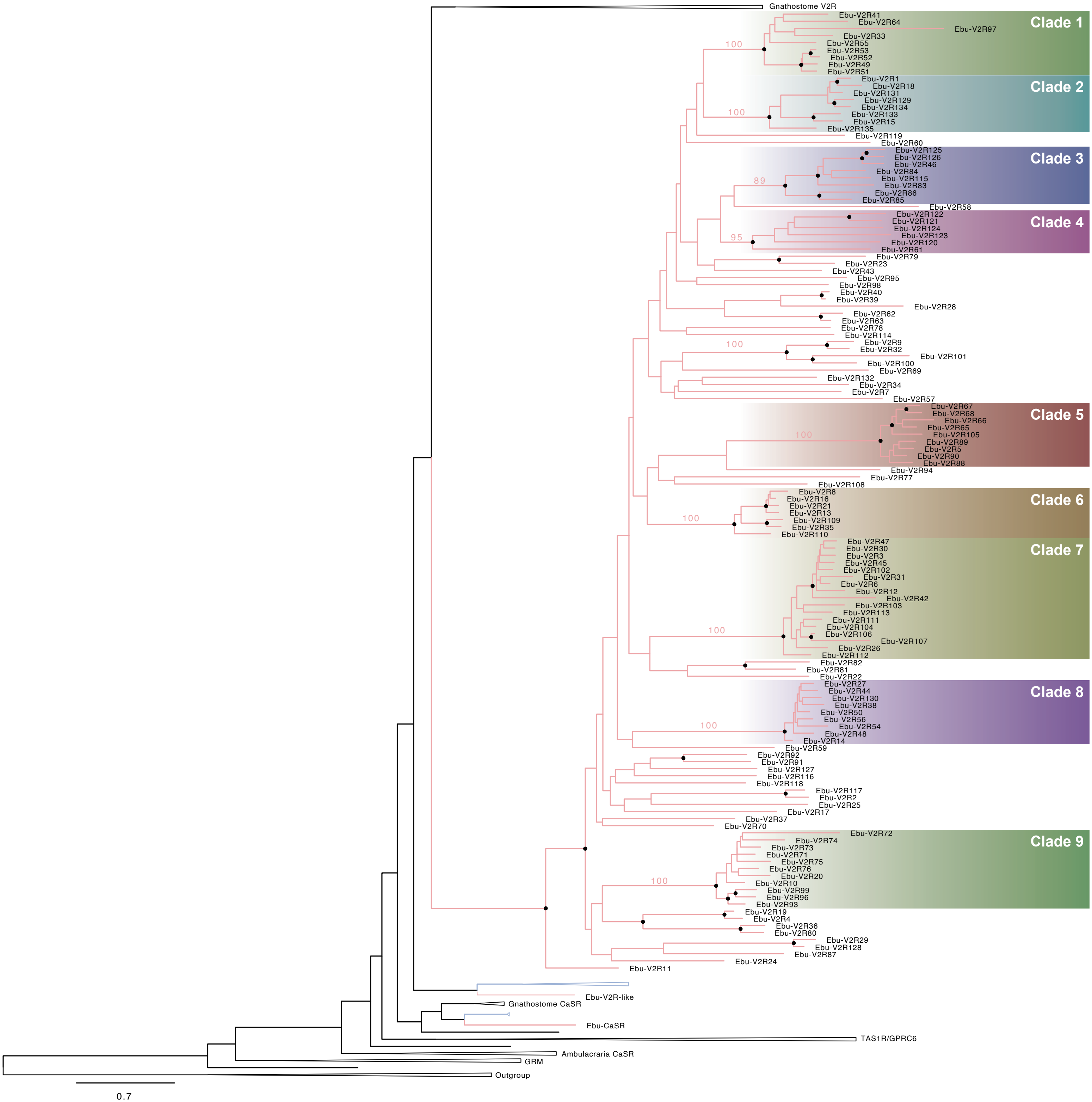

### Supplemental Fig. 3

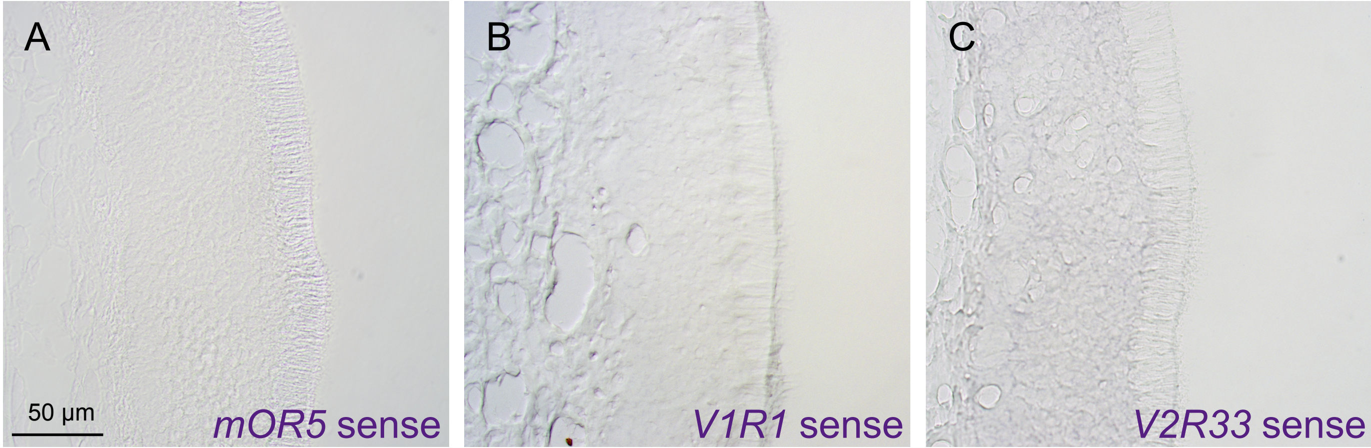

### Supplemental File 4

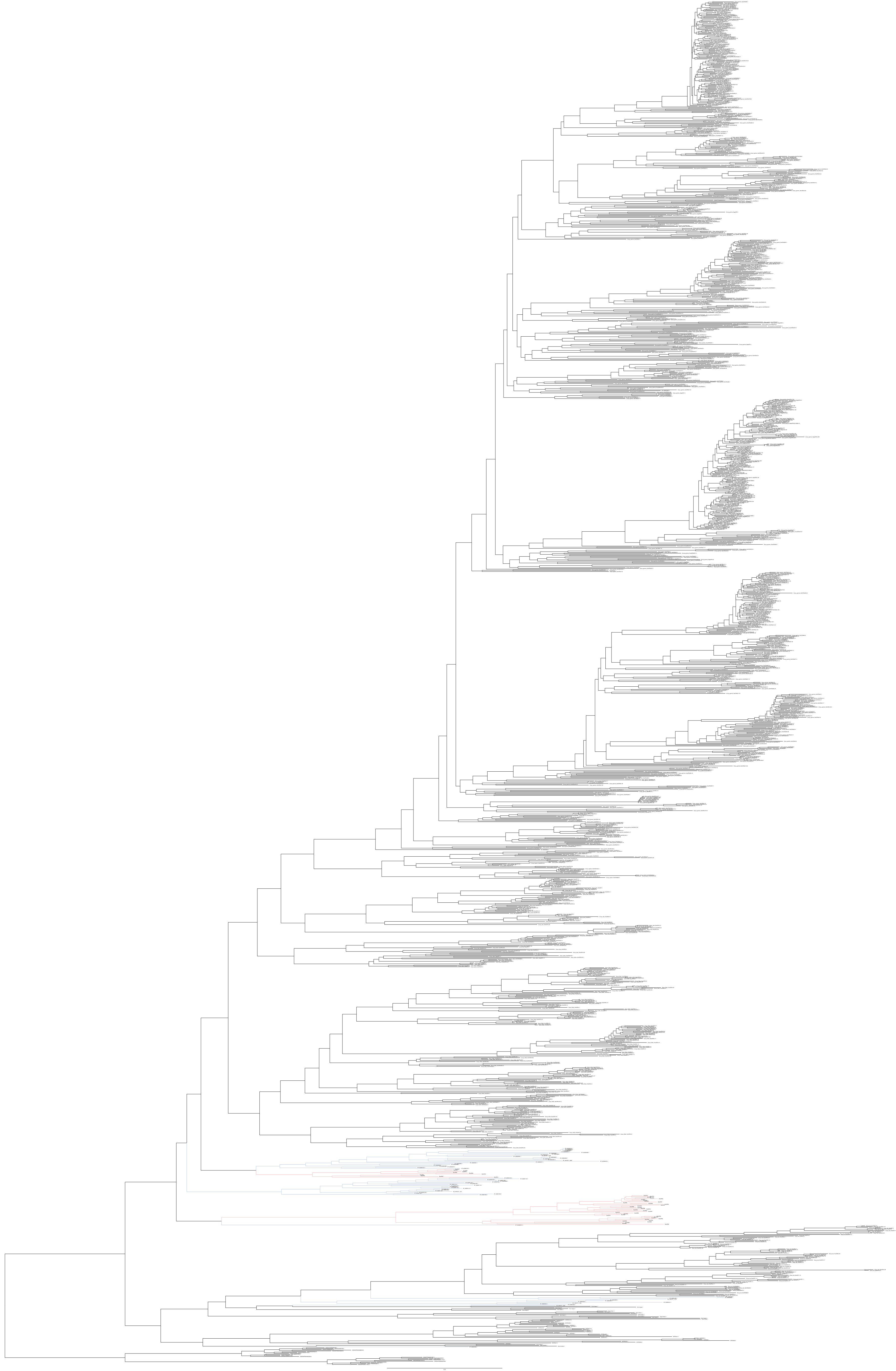

### Supplemental File 5

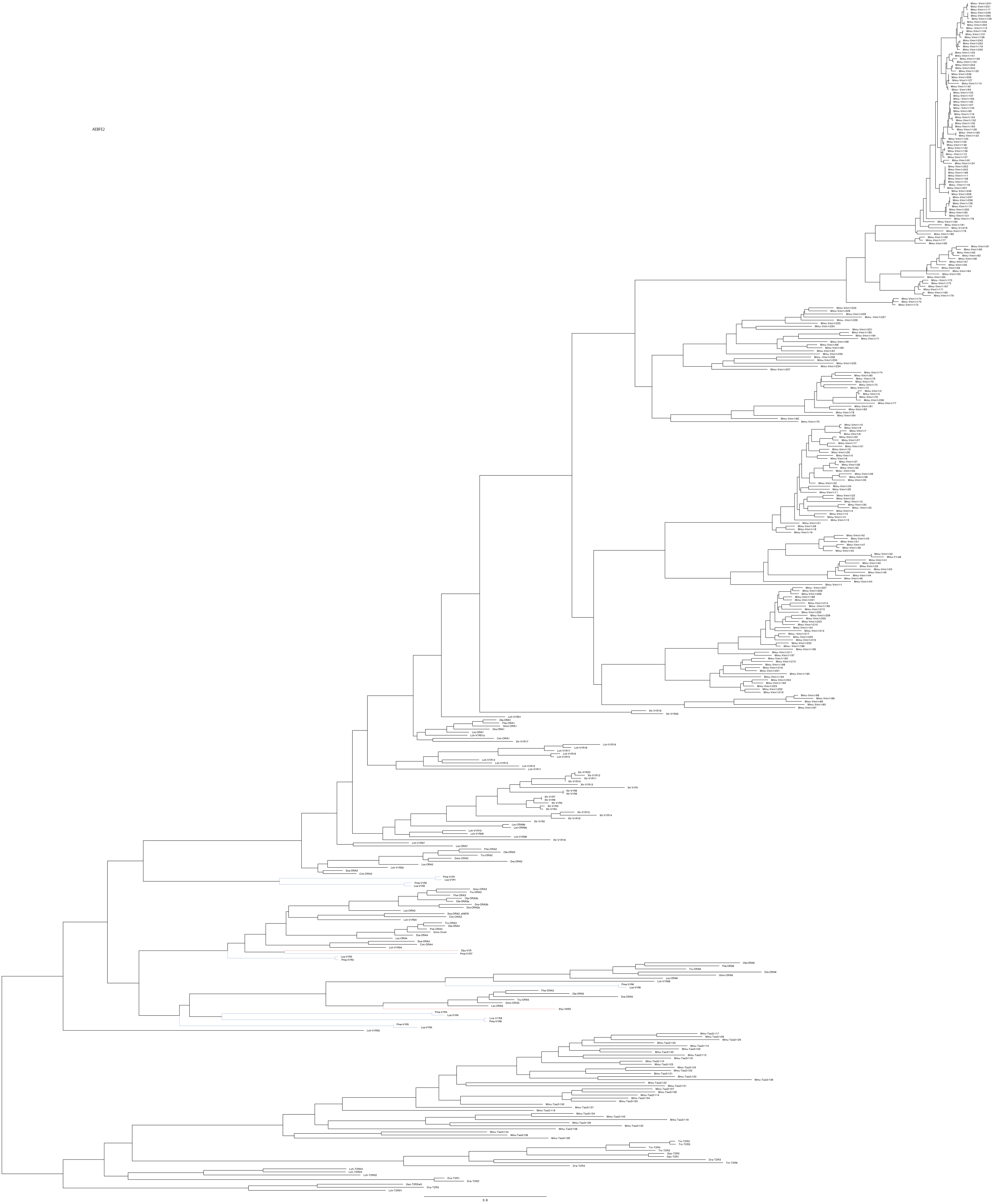

### Supplemental File 6

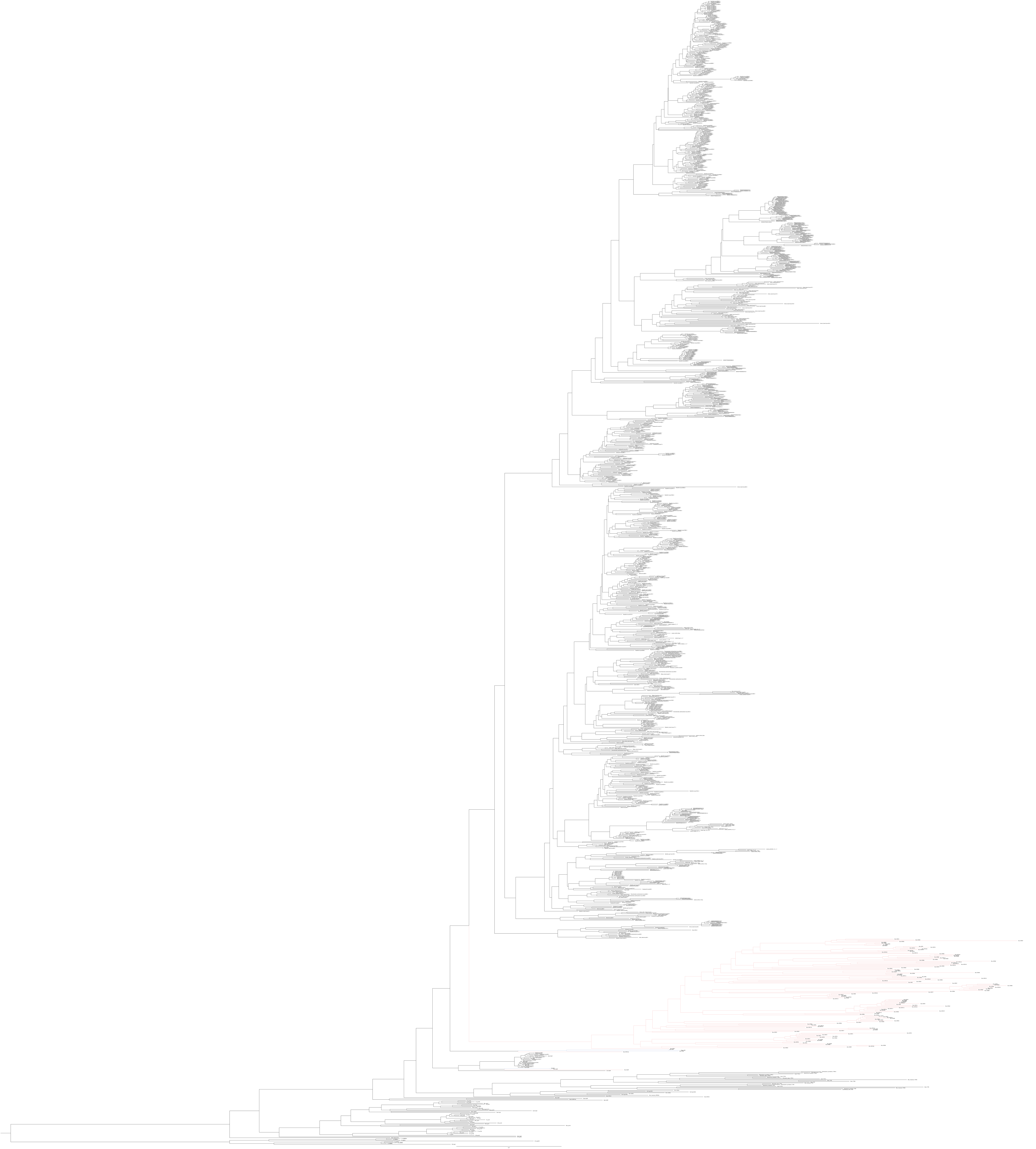

### Supplemental File 7

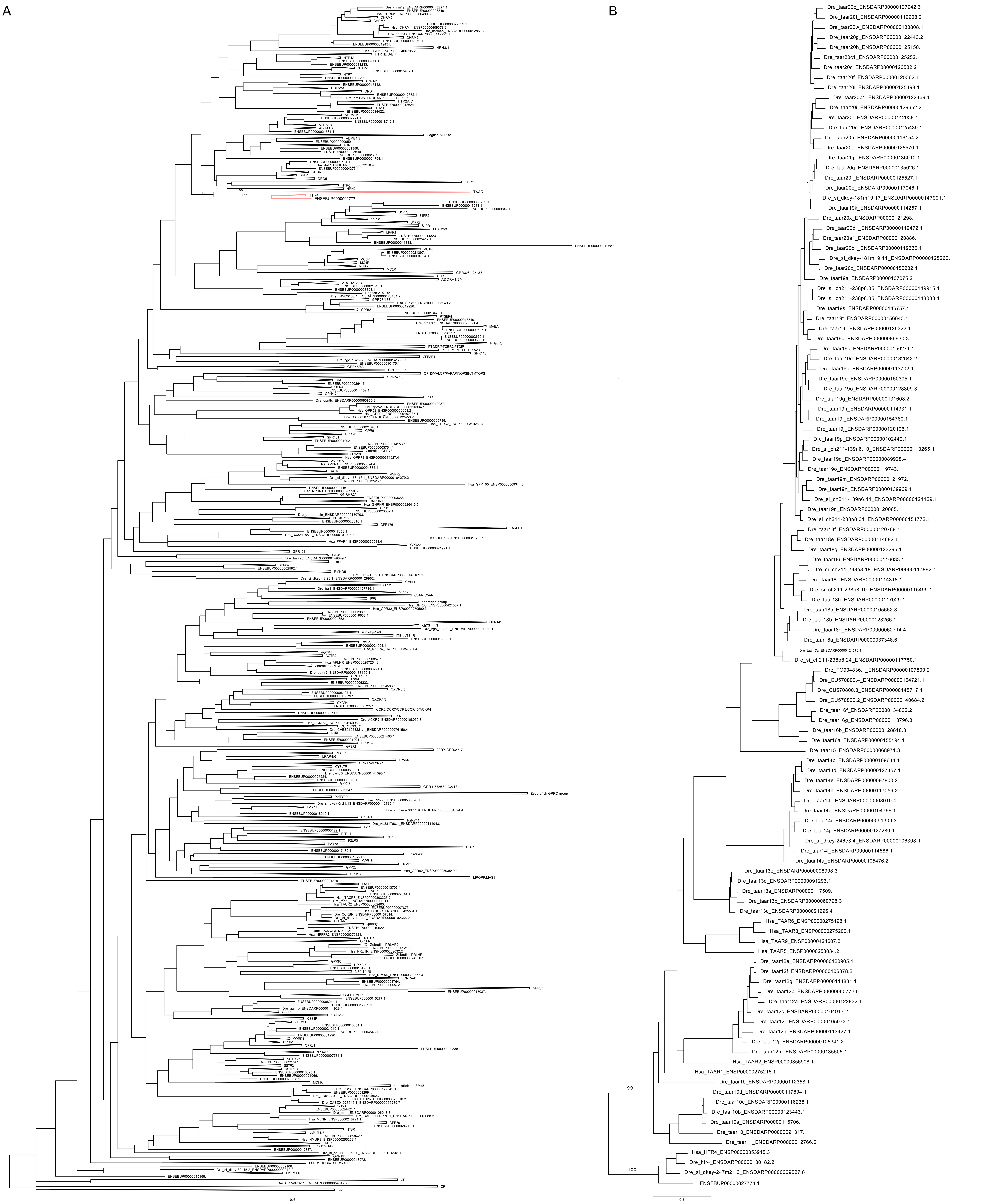
